# Identifying gene regulatory networks of senescence in postharvest broccoli (*Brassica oleracea*)

**DOI:** 10.1101/2021.02.18.431815

**Authors:** Yogesh Ahlawat, Song Li, Eleni D. Pliakoni, Jeffrey Brecht, Tie Liu

## Abstract

The facts of postharvest food loss and waste and the resulting consequences affect us in many ways, ranging from important economic and social issues to lasting and detrimental environmental problems. We are using genomic tools to understand senescence in postharvest broccoli florets when stored at room temperature or 4 °C. The RNA-sequencing approaches provide key insights into the gradual changes in transcriptome profiling in broccoli during postharvest storage. Identification of those key factors could lead to a better understanding of gene regulation of postharvest senescence. Those genes could serve as ‘freshness-indicators’ that have the potential to mediate senescence and to generate germplasm for breeding new varieties with longer shelf-life in Brassica vegetables. Such a tool would also allow a new level of postharvest logistics based on physiological age, supporting improved availability of high-quality, nutritious, fresh vegetables and fruits.

**One sentence summary:** A gene regulatory network modulates senescence in postharvest broccoli florets

## Introduction

With the exception of a few climacteric fruits that may ripen postharvest, fresh fruits and vegetables begin to lose quality immediately after harvest and during processing, transportation, and storage. Produce freshness is an important and fleeting state usually associated with proximity to harvest and consumer desirability, but otherwise it is not well defined (Peneau et al., 2006; Ragaert et al., 2004). Tissue physiological age (Ashby, 1950) may more precisely define freshness as opposed to chronological age or time after harvest. During the postharvest period, physiological age is intimately related to the process of senescence, which is initiated by removal of the organ from the plant, arguably making at-harvest freshness the initial stage of senescence for detached plant organs. Senescence is defined as the series of “processes that follow physiological maturity or horticultural maturity (i.e., harvest maturity) and lead to death of the tissue” (Watada et al. 1984). Therefore, freshness, physiological age, and senescence are complementary concepts – freshness indicators decline as physiological age and senescence advance (Schwerdtfeger, 1979). With increasing physiological age and progression through the different stages of senescence come changes in gene expression, protein synthesis, physiological processes, and various physical and compositional markers (Humbeck 2014; Nooden et al., 1997).

Using broccoli (*Brassica oleracea* L.) and Romaine lettuce (*Lactuca sativa* L. var. *longifolia*) as test crops, we measured color, weight loss, chlorophyll *a* and total chlorophyll content, chlorophyll fluorescence, phenolic content and polyphenol oxidase (PPO) activity, vitamin C content, and protein content in order to evaluate the usefulness of those factors as senescence indicators and predictors of remaining postharvest life (Brecht et al., 2014; Pliakoni et al. 2015). However, we concluded that the physiological indicators of senescence measured in isolation as “snapshots” rather than followed over time are not useful for determining physiological age or freshness because their levels vary widely from lot to lot and even among different harvests from the same plants. Lack of reliable indices for defining produce freshness limits our capacity to control produce quality and results in huge amounts of food loss and waste. What is needed to assay the physiological age or stage of senescence of plant tissues are reliable, reproducible, and easily assayable markers - in other words, compounds that are either present or absent at different stages of senescence.

### Senescence associated genes (SAGs)

During senescence, several processes occur: deterioration of cell structure, decline of photosynthesis, and degradation of macromolecules such as proteins, nucleic acids, and lipids. Biochemical and molecular studies have shown that senescence is an active process that involves expression of many new genes and synthesis and modification of proteins (Nooden et al., 1997). Many of our recent insights into plant senescence have come from research with the model plant, *Arabidopsis thaliana* (The Arabidopsis Genome Initiative, 2000; Breeze et al., 2011; Woo et al., 2010), extended to other cruciferous plants such as broccoli (Gapper et al., 2005; Gonzalez and Botella 2003). These genes were designated as SAGs, and include those related to chlorophyll degradation, hormone response pathways, protein degradation, and lipid/carbohydrate metabolism (Li et al. 2012). The Leaf Senescence Database (http://psd.cbi.pku.edu.cn/) covering genes associated with leaf senescence in 44 plant species, including several Brassica species, is a key resource for researchers in plant senescence.

Postharvest changes in immature broccoli florets have shown many similarities to molecular and biochemical changes occurring during leaf senescence (Page et al. 2001). In broccoli florets, according to Chen et al. (2008), the senescence-associated genes include ethylene-related genes in the ACO (*BoACO1, BOACO2, BoACO3*) and ACS (*BoACS1, BoACS2 and BoACS3*) gene families, metabolic-related genes like chlorophyllase (*BoCLH1, BoCLH2, and BoCLC3*), and other functional genes like cysteine proteases *(BoCP5, SAG12, LSC790, and LSC7*), and glutamine synthetase (*LSC460*). Also, Eason et al. (2014) reported that protease activity is involved in the postharvest senescence of detached broccoli. By overexpressing the *BoCPI-1* inhibitor in broccoli, they observed a delay of chlorophyll loss after harvest.

It has been reported that during the progression of senescence, certain SAGs are upregulated or downregulated. With regard to the ethylene-related genes, it has been reported that some are barely detectable in freshly harvested immature florets (*BoACO2 and BoACO3*), while others like *BoACO3* are only expressed in whole yellow florets (Ahlawat and Liu, 2021). There are also other examples of broccoli genes that exhibit different expression patterns after harvest and through senescence; for example, *BoACS1* (DNA clone of ACC synthase) had high transcript levels after harvest but was not detected during storage. Also, *BoACS2* increased at the last stages of senescence, but it was stable until then, and *BoACS3* was not detectable until the final stages (Gonzalez and Botella 2003). Additionally, three chlorophyllases have been cloned, and only the transcript of chlorophyllase *BoCLH1* is visible in freshly harvested and 4-day postharvest florets in Northern blot analysis, while mRNAs of *BoCLH2* and *BoCLH3* are barely detectable (Chou, 2001). *BoCAB1*, a putative chlorophyll a/b-binding protein, the major component of light-harvesting, declined rapidly after harvest (Gapper et al., 2005).

To maintain the high levels of respiration, the transcripts of genes involved in sucrose metabolisms, such as BoINV1 and BoHK1, and in sucrose transport, including BoSUC1 and BoSUC2, are upregulated during senescence (Coupe et al. l., 2003; Gapper et al., 2005). The genes involved in protein degradation and in the amino acids recycled from degraded protein are thought to be upregulated in senescing tissues. In harvested broccoli, the transcripts of several protease genes, such as cysteine proteases *(BoCP5, SAG12, LSC790, and LSC7*), and glutamine synthetase (*LSC460*), the action of which involves the conversion of free amino acids released from degraded protein into glutamine to transport, are increased (Eason et al., 2005; Gapper et al., 2005; Kamachi et al., 1991; Page et al., 2001). Jara et al. (2019) characterized the expression of a gene encoding a chlorophyll b reductase (*BoNYC1*), a critical enzyme in chlorophyll catabolism, during postharvest senescence of broccoli. They found a close correlation between chlorophyll degradation and expression of the *BoNYC1*.

In this study, using broccoli floret tissue, we take the first steps to test the hypothesis that aging of produce (i.e., loss of freshness) after harvest may be divided into discrete episodes of senescence processes that are defined by diagnostic patterns of gene expression profile changes.

## Results

### Weight loss and chlorophyll degradation in postharvest broccoli

To evaluate the physiological changes during postharvest senescence in broccoli, we measured weight loss and chlorophyll content. Monitoring the rate of weight loss in a 10-day period of time showed that there is significant decline in weight within the first 3 days of storage. The weight decreased by 18% and 14% in room temperature and cold storage, respectively, although there was no visible yellowing and chlorophyll decline observed during this early stage of postharvest storage (Fig. 1A-C). We observed the florets yellowing in broccoli after 5 days of postharvest storage at room temperature (∼25 °C) in contrast to the tissue stored at a refrigerator temperature of 4 °C (Fig. 1C). To quantitatively evaluate the color changes in broccoli, we measured the total chlorophyll content (Fig. 1C). In both conditions, a general progressive decrease of chlorophyll concentration was observed from day 3 on, although no obvious color changes were found in the early storage time period. However, chlorophyll content was reduced significantly in room temperature storage (Fig. 1C). Based on the changes in weight loss and chlorophyll content during postharvest storage, we predict that there are rapid physiological and biochemical changes in early storage of broccoli.

**Figure 1.**
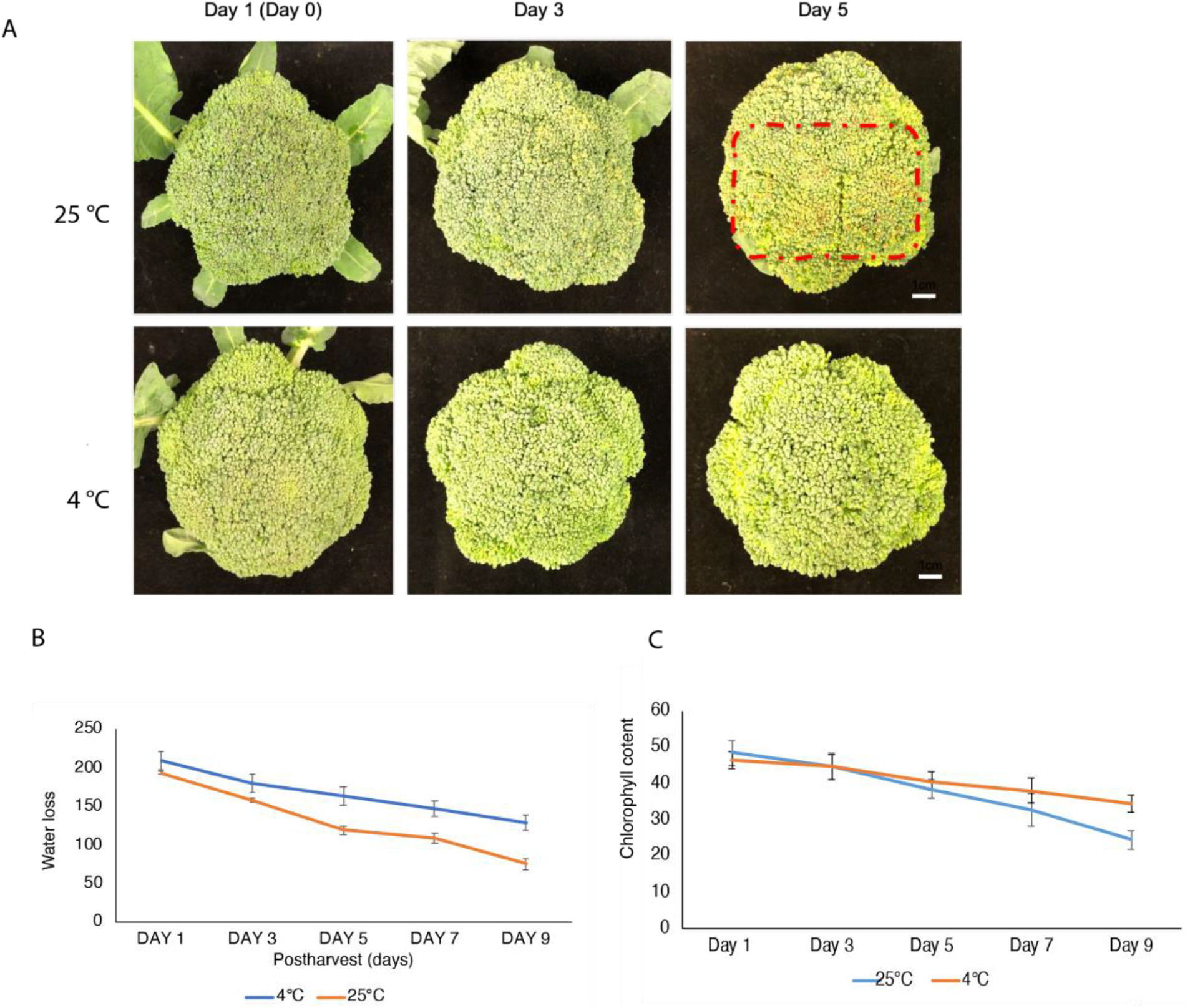
Broccoli heads at 5 days room temperature and cold storage. A. Appearance of broccoli heads. B. Water loss in postharvest broccoli. C. Chlorophyll content decrease caused by postharvest senescence in broccoli.

### Dynamic regulation of differentially expressed genes across different conditions

To investigate the rapid response to postharvest stresses, we generated a time-course transcriptome of broccoli crown at three time points (immediately after harvesting, 3 days, and 5 days postharvest) and two storage conditions (room temperature ∼25 °C and 4 °C cold storage) using RNA-sequencing (RNA-seq). Three biological replicates for each time point and storage condition were used. Principal Component Analysis (PCA) was performed to detect the similarity and variability among all the samples. The plot showed that samples at later time points are well separated from the first time point (1 day). The first and second principal components represent the changes between the 4 °C treatment or 25 °C at 3 and 5 days, respectively. The biological replicates are grouped together along the second component (Fig. 2A), suggesting much smaller variations between biological replications as compared to the experimental treatments. We analyzed transcripts whose expression was significantly changed between the two storage conditions at each time point and between different dates (Fig. 2B-C). We have found a number of differentially expressed (DE) genes that varied depending on those time points. When we compared the gene expression under the two different storage conditions, we found 4279 transcripts were DE within the first 3 days (2768 up-regulated and 1511 down-regulated), implying that there is a rapid gene induction in the early postharvest senescence stage. In addition, we found 4143 transcripts were DE at the fifth day of storage (1503 up-regulated and 2640 down-regulated; Fig. 2B).

**Figure 2.**
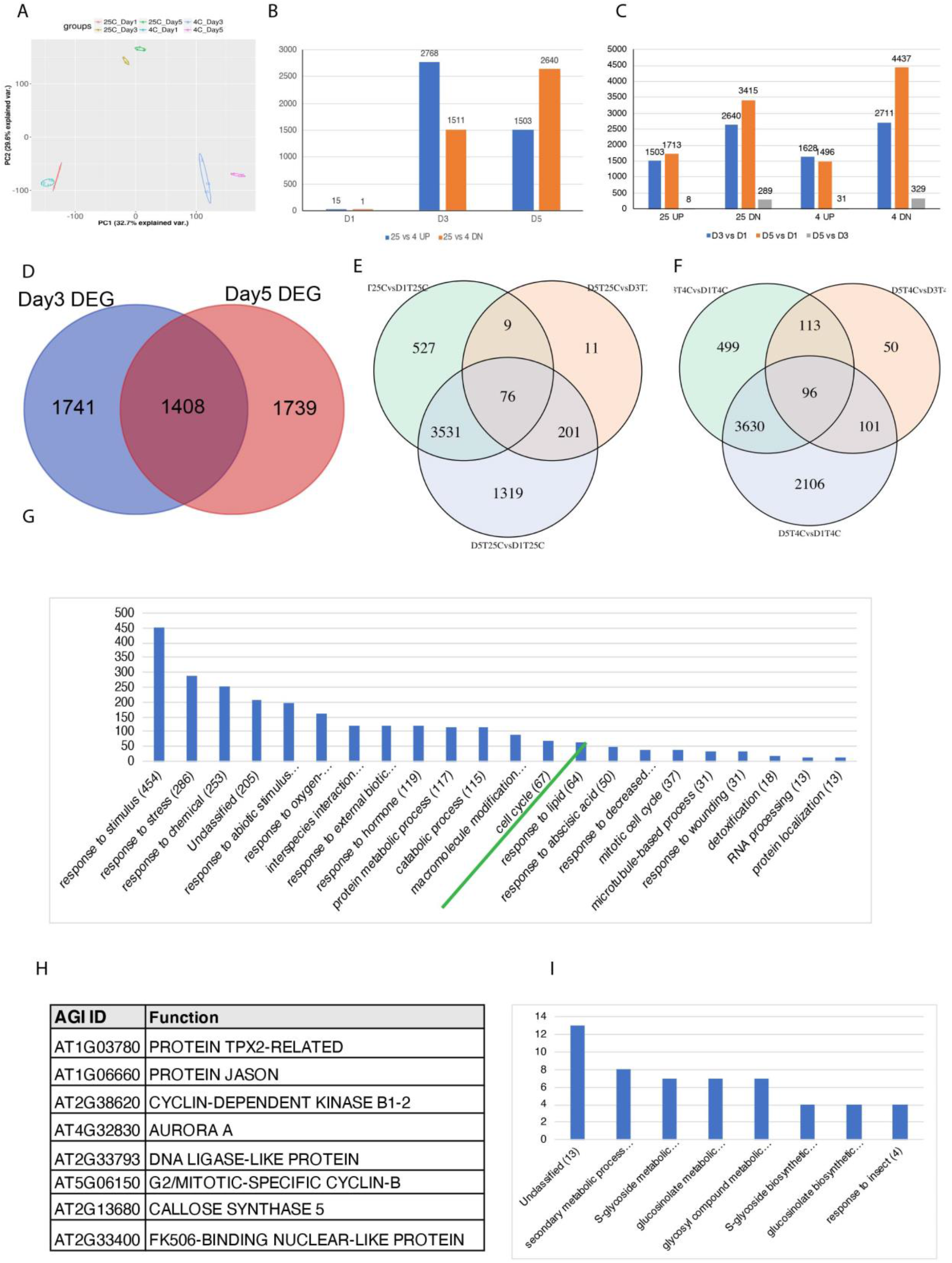
Identification of differentially expressed genes (DEG) through RNA-seq. A. Principal-components plots of RNA-seq data. Color code is shown. Each dot represents one sample. Color-coded cycles show the replicates. B. Total number of significantly differentially abundant (>2 folder changes) genes in each time points detected in comparison between room temperature and cold storage. Blue represents the number of upregulated genes; red represents that of down-regulated genes. C. Total number of DEGs (>2 fold) induced and repressed genes detected in room temperature and cold storage condition. D. Hierarchical clustering of relative expression (RE) of detected genes at each time point; E. Under room temperature and cold storage. Ratio calculated as log2.

Interestingly, the induced and repressed genes displayed opposite expression patterns at 3 days versus 5 days. Whereas the number of induced genes decreased by 46%, from 2768 to 1503, the number of repressed genes significantly increased by 42%, from 1511 to 2640. This led us to speculate that the early postharvest response is predominantly a rapid induction of gene expression, which probably reflects stress response during the early postharvest stage. In addition, many more genes were repressed beginning at the 5 days’ time, suggesting that various biological processes slow down when broccoli is stored at room temperature. We next compared the up-regulated and down-regulated DE genes at room temperature and cold storage across the time courses analysis. Results of these comparisons are shown in Fig. 2C. As can be seen, the number of upregulated genes at room temperature and cold storage were similar, as well as the number of downregulated genes, except that the number of down-regulated genes on the fifth day of cold storage was much higher than at room temperature, thus illustrating the well-known fact that refrigerated storage slows the negative, senescence-related changes in postharvest produce (Lauxmann et al. 2012).

To test our hypothesis, we next performed Venn Diagram and Gene Ontology (GO) enrichment analysis on the DE transcripts under the two storage conditions in the early (3 days) and late (5 days) postharvest timepoints. In comparing the two storage conditions, we found 1408 DE genes (25 °C/4 °C) were consistently expressed at both timepoints (Fig. 2D). GO enrichment analysis showed that those genes were primarily related to the ‘cellular process’, ‘response to stimulus’, and ‘response to abiotic and biotic stress including wounding and osmotic stresses’ categories, suggesting that various stress responsive pathways were functioning in postharvest broccoli across different conditions at the mRNA level (Fig. 2G). In contrast, when we examined the overlapping DE transcripts among all timepoint groups (D3/D1, D5/D1, and D5/D3), we identified 76 DE genes, but only under room temperature conditions (Fig. 2E). Interestingly, GO analysis showed that a number of cell cycles were enriched in those overlapping gene groups (Fig. 2H), indicating that senescing cells may represent a transient phase similar to a de-differentiation event, which involves either a re-entry into cell cycle or a trigger to cell death. This is consistent with the previous discovery that senescing cells may share common features with de-differentiating cells (Damri, et al. 2009). We next identified the overlapped DE genes only in cold storage and found 96 genes that were shared among all DE timepoints groups (Fig. 2F). Specifically, GO enrichment analysis showed that genes related to glucosinolates and secondary metabolite pathways are highly expressed in the overlapped set of genes during the cold condition (Fig. 2I). The expression of the glucosinolates and secondary metabolic pathway genes were conserved in all timepoints during cold storage, suggesting that there are nutritional remobilization events during tissue senescence in postharvest broccoli. These analyses revealed that the defense-growth trade-off pathways are consistently activated during postharvest senescence even when broccoli was held in refrigerated conditions.

### Time-specific gene expression during postharvest senescence in broccoli

To address what might specify differences among gene transcripts that change during the stages of postharvest senescence, we identified putative time-specific genes at two given time points (3 days and 5 days). Comparing the number of DE genes between room temperature and cold treatment (25 °C/4 °C), we observed 1827 and 791 genes that were specifically expressed in room temperature at the day 3 and day 5 timepoints, respectively (Fig. 3A). We found 635 genes that are overlapped between those two datasets (Fig. 3A, Supplementary Table 1). Besides the genes involved in response to abiotic and biotic stimulus, GO enrichment analysis showed genes related to senescence, aging, and response to oxidative stress among the overlapped genes (Fig. 3C). Additionally, as observed for DE transcripts on day 3 in Fig. 3C and Fig. 4A, 40 genes in the “response to chitin” group and 41 genes related to organonitrogen compounds were highly expressed in the room temperature condition. An organonitrogen compound is formally a compound containing at least one carbon-nitrogen bond. We also found that genes for ‘response to wounding’ and ‘response to oxygen-containing compound’ were DE in the early senescence stage (Fig. 4A, highlighted in the orange and red boxes), suggesting defense-related genes are specifically expressed at room temperature on those timepoints.

**Figure 3.**
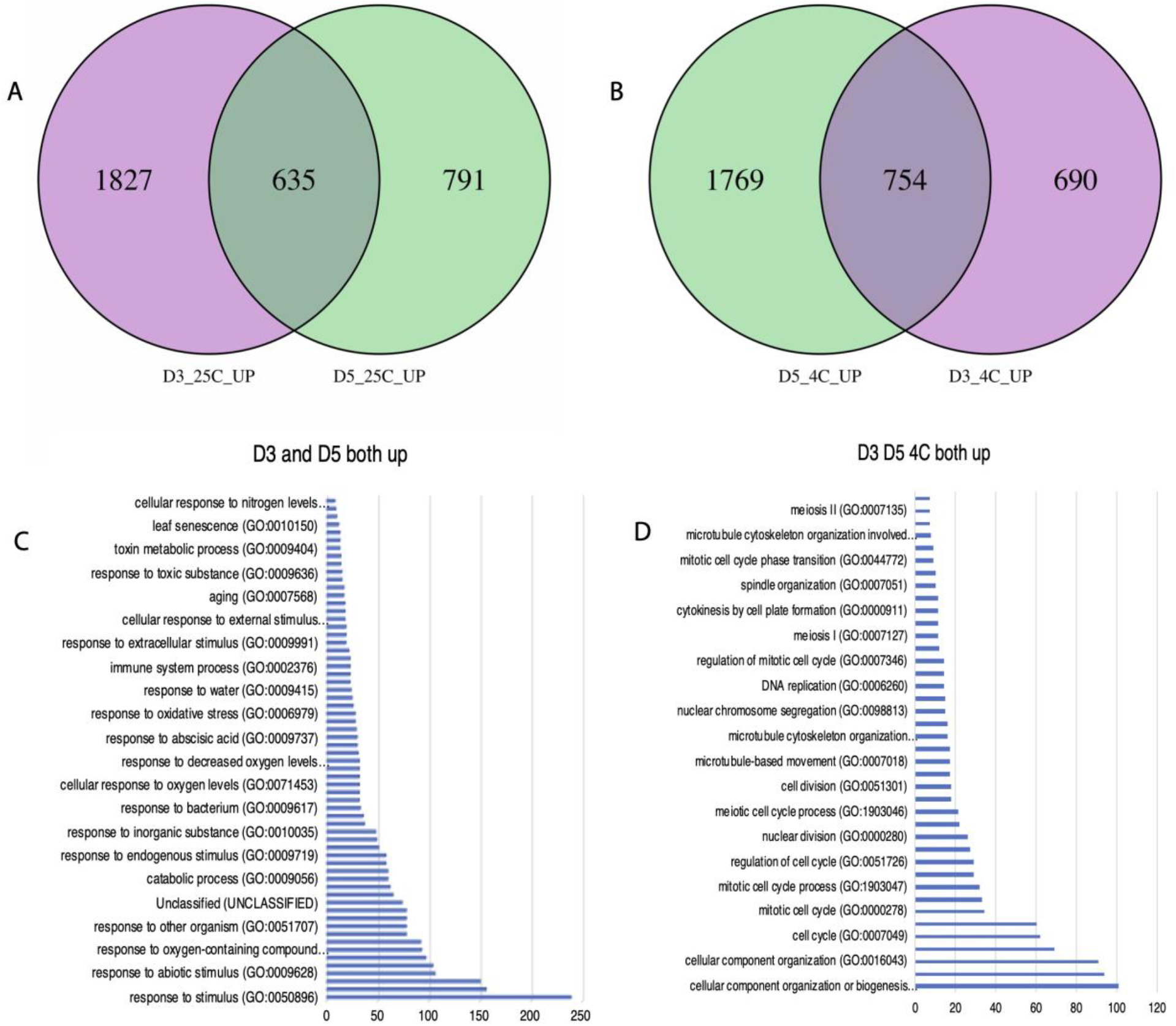
Overview of overlapped genes between room temperature and cold storage. A-B. Venn diagram illustrating the overlap DEGs between day 5 and day 3 under room temperature condition; and cold storage condition, B. C-D. A list of GO terms enriched in the clusters is shown for overlapped DEGs at room temperature, C; in cold storage, D.

**Figure 4.**
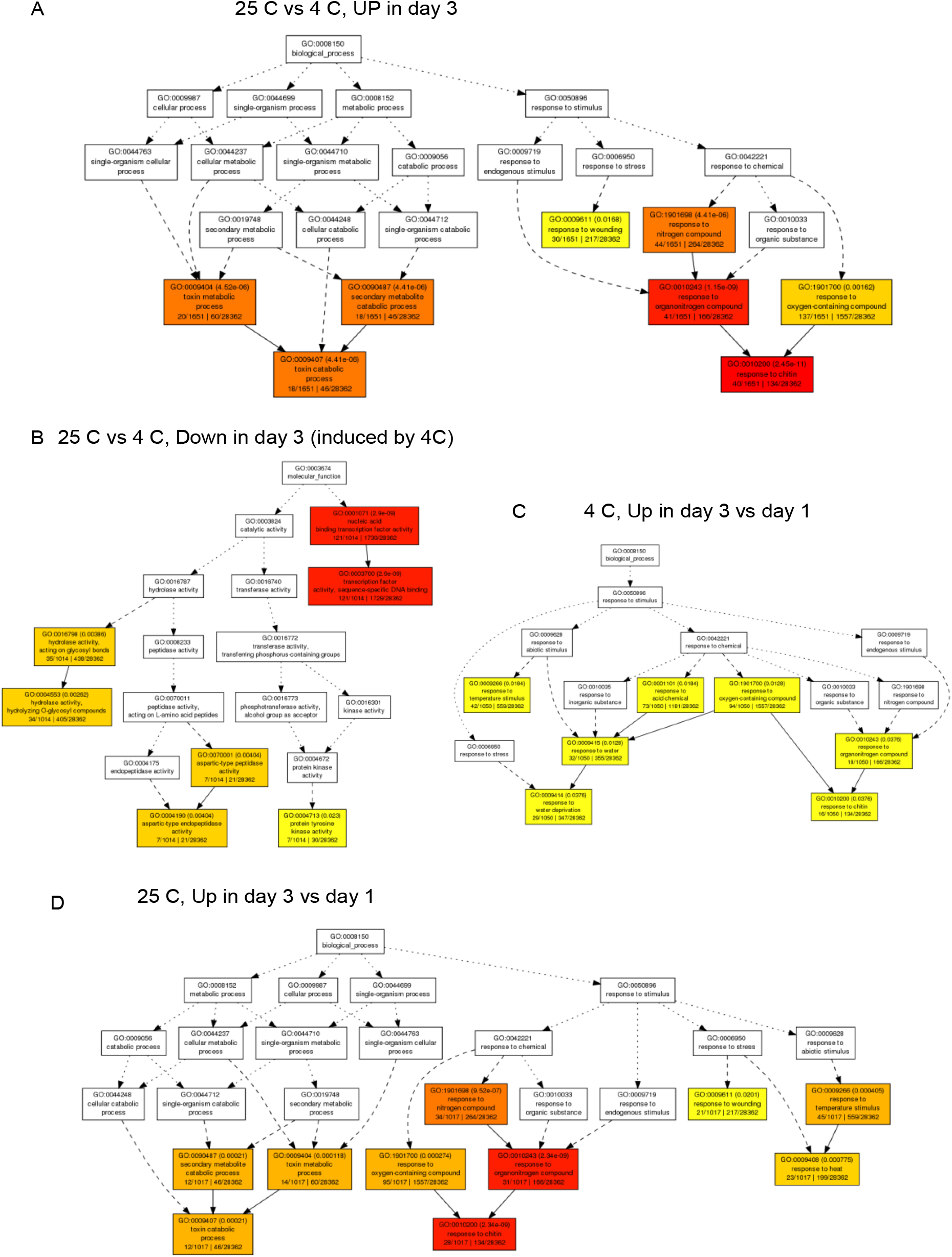
A list of GO terms analysis of DEGs. A. The upregulated genes at 25 °C versus 4 °C. The down-regulated genes at 25 °C versus 4 °C. C. The upregulated genes at room temperature (25 °C) on day 3 versus day 1; D. The upregulated genes on day 3 versus day 1 during cold storage (4 °C). Color is related to the significance p value. The red ones are more significant than orange and then yellow. The p values are provided in each box (for example, GO:0009407, p value is 4.41e-06). Number of genes in the list and in the genome are also provided in the box.

On the other hand, 121 genes related to ‘nucleic acid binding transcription factor activities’ were highly expressed in the cold treatment at the day 3 and day 5 (Fig. 3B). For the genes specifically expressed on the day 5 timepoints, we found 791 genes at room temperature and 1769 genes in the cold treatment (Fig. 3B). GO enrichment analysis showed that genes related to ‘regulation of gene expression’ and ‘response to chemicals’ were highly expressed on those timepoints in room temperature storage. In contrast, we found ‘developmental process’, ‘cell cycle’, and ‘cell wall associated’ genes were specifically expressed in the cold treatment on the day 5 (Fig. 3D and 4C). Collectively, genome-wide comparisons between different storage conditions and timepoints analysis illustrate the temporal and spatial-specific genes. Those potential candidate genes could also serve as markers to monitor the senescence stages of postharvest broccoli.

### Validation of RNA-seq data through Real-Time PCR and transient tobacco analysis

To validate our DE results, we used three criteria for selecting candidate genes within the networks for further analysis: the level of expression, functional category, and how ‘connected’. We set up three experiments to test the top candidate genes. We first measured the transcript levels of 10 selected genes under both storage conditions using Real-Time PCR (qPCR). A side-by-side comparison between RNA-seq and qPCR is illustrated in Fig. 5. Based on the functional category, we selected two SAGs, *SAG2* (*LOC10637874*) and *SAG12* (*LOC10632502*); two genes encoding peroxidase (LOC106325226) and (LOC106325226); two stress-responsive genes, *MYB49* (*LOC106332869*) and *GT61* (Glycosyltransferase family 61 protein, *LOC106343191*); two growth regulators, dormancy/auxin-associated protein (*LOC106317320*) and IDA (*LOC106299577*); and two cell-wall associated genes, methyltransferases (*LOC106301029*) and pectin transferases (*LOC106325531*). As shown in Fig. 5, the RNA-sequencing and qPCR results displayed very similar patterns. For instance, the transcript levels of both *SAG2* and *SAG12* rapidly increased in response to postharvest stresses, with over 600-fold and 30-fold changes, respectively.

**Figure 5.**
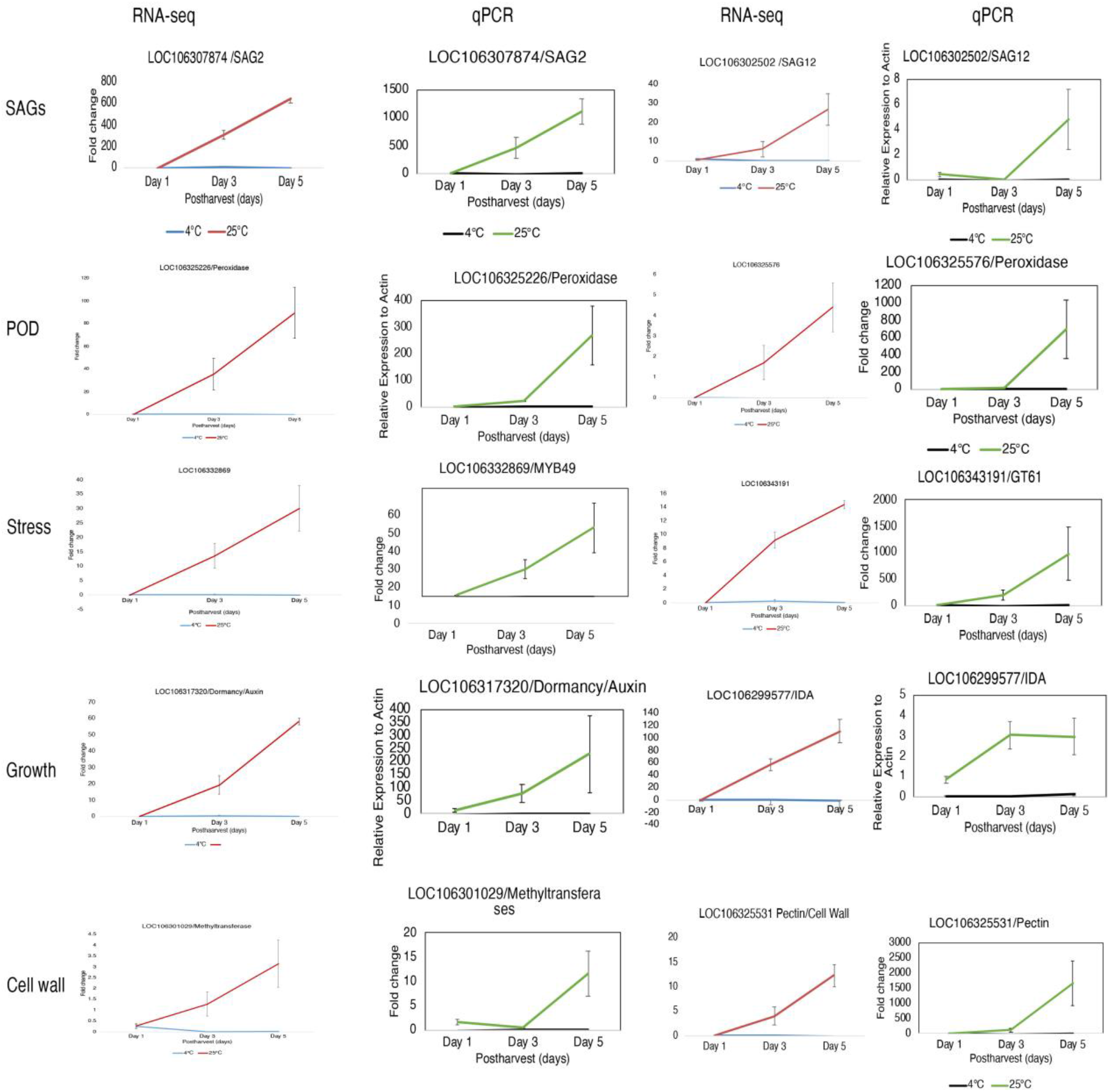
Comparison of RNA-sequencing and real-time PCR analysis for the top DEGs. The relative expression levels are generated from fold changes. The error bars indicate the standard error. A-D. Senescence-associated genes. E-H. developmental factors; I-L. Reactive oxygen species (ROS); M-P. stress-induced genes; Q-T. cell-wall-related genes.

To further validate the results, we selected eight genes from a short list of 45 top DEGs that were rapidly induced on day 3 at room temperature in comparison to the cold storage condition. The genes are listed in Fig. 6A. To test if those genes play a role in tissue senescence, we overexpressed those genes and examined phenotype through transgenic tobacco plants. We postulated that overexpression of those genes could trigger senescence that led to accelerated tissue deterioration. We performed transient assay in tobacco epidermal tissue to visualize progression of tissue senescence (Fig. 6B-D, and Supplementary Fig 1). As expected, in a negative control we didn’t observe any phenotypic changes of transgenic tobacco plants expressing empty vector (pMDC43) after 3 and 5 days post-injection. Interestingly, we found tissue yellowing and browning in the overexpresser of *ZPR3* (*pMDC43:ZPR3*) and *DUF642* (*pMDC43:DUF642*) at 3 days post-injection (Fig. 5C-D). More obvious color bleaching and yellowing were observed at 5 days post-injection. Similar results were found in overexpression of *NAC46* and *S-adenosyl-L-methionine-dependent methyltransferases (S-ALM)* (Supplementary Fig 1). These findings confirm that those DEGs have a potential role in tissue senescence.

**Figure 6.**
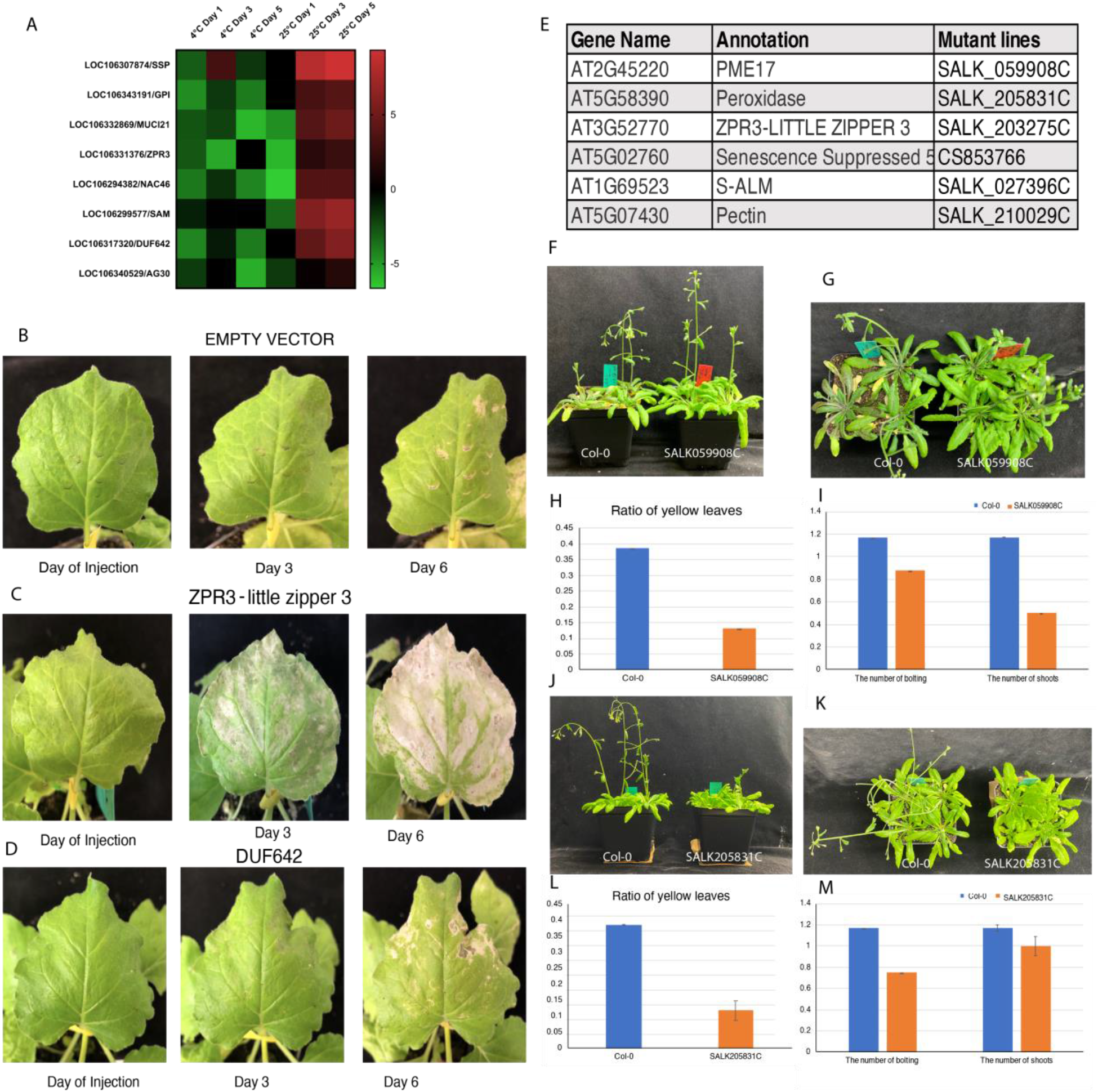
Validation of genes involved in senescence through tobacco transient assay and *Arabidopsis* mutant analysis. **A**. A heatmap of selected genes for tobacco transient assay. B. Representative images of control tobacco leaves injected with pMDC43 empty vector. C. Overexpressing *ZPR3* (*pMDC43::ZPR3*) in tobacco leaves caused tissue browning and accelerated tissue senescence. D. Overexpressing of *DUF642* (*pMDC43::DUF642)* in tobacco leaves displayed progressive senescence after injection. E. Selected DEGs of *Arabidopsis* homologues and their T-DNA insertion mutants. F-G. *SALK059908C*, a loss-of-function of PME17 showed delayed maturation and senescence phenotypes. H. Number of yellow leaves in total leaves in *SALK059908C* mutants (n=8). I. The number of bolting and the number of shoots were scored in wildtype and mutant. J-K. *SALK205831C*, a loss-of-function of peroxidase gene showed delayed senescence. L-M. Number of yellow leaves in total number of leaves (n=8). M. The number of bolting and the number of shoots in wildtype and *SALK205831C* mutants.

In a third validation experiment, we took advantage of the genetic resources of *Arabidopsis* mutant collections and identified the broccoli homologous genes in *Arabidopsis*. Ten *Arabidopsis* mutants from a short list of 45 top DEGs were obtained from the Arabidopsis Biological Resource Center (ABRC) stock center (The Arabidopsis Information Resource, TAIR). We selected those T-DNA insertion lines that contain insertional mutation in the exon and are homozygotes in *Arabidopsis* (O’Malley, et al. 2017). The genes and insertional mutants were listed in Fig 6E. We grew those mutants in the same flat with wildtype Col-0 control and scored the mutant phenotype once bolting occurred. We quantified the phenotypes of mutants by comparing the ratio of yellow leaves to total leaves, the number of axillary shoots, and the number of bolting (i.e., flowering) shoots. We observed 5 out of 10 mutant lines displayed a smaller number of yellow leaves and a lower number of axillary shoots and bolting shoots (Fig. 6F-M, and Supplementary Fig 2). We note that this is a quantitative observation to confirm that loss-of-function of those mutants resulted in delay of senescence. Two of the mutant lines (*SALK059908C* and *SALK205831C*) with their quantitative data are illustrated in Fig. 5F-M. These results suggest that the *Pectin Methylesterase 17* (*PET17*) and *peroxidase family gene (At5g58390)* loss-of-function mutants exhibit delayed leaf senescence. Additionally, loss-of-function mutants of *S-adenosyl-L-methionine-dependent methyltransferases (S-ALM, At1g69523)*, pectin lyase-like family gene (At5g07430), and *zpr3* displayed similar phenotypes that delayed senescence (Supplementary Fig 2).

### Comparative analysis of Arabidopsis and broccoli to disclose a conserved senescence gene-regulatory network (GRN)

To investigate the specific genes in broccoli relevant to postharvest senescence, we utilized the existing dataset from functional genomics studies of senescence in *Arabidopsis* (Breeze, 2011; Su, 2013). When we merged our day 3 and day 5 DEGs, we found that a total of 4279 genes were differentially expressed in the room temperature condition on either of the timepoints. We identified the Arabidopsis homologues for those 4279 DEGs and conducted comparative analysis of DEGs among postharvest broccoli, leaf senescence in *Arabidopsis*, and flower senescence in *Arabidopsis*. Breeze et al (2011) identified 6323 DEGs through a time-course profile of microarray analysis in a single *Arabidopsis* leaf over a 3-week period of senescence. We reasoned that investigating the regulatory network governing leaf senescence would help deepen our understanding of the common molecular cellular mechanism during tissue senescence across various tissue types. To minimize biases introduced by potential leaf-specific expression of senescence, we further compared the 4153 genes that were identified from a transcriptional study on stress-induced flower senescence in *Arabidopsis* (Su, 2013). This study helps to identify tissue-specific genes during senescence. The combination of three genomics datasets may help us define the common and tissue-specific SAGs.

Venn diagram analysis revealed that there were 429 commonly expressed DEGs (ceDEGs) in all three combinations of the dataset. Those genes were either up-regulated or down-regulated (Fig. 7A) and are listed in Supplementary Table 1. Gene Ontology enrichment analysis of the 429 overlapped genes showed that the majority of the genes in this group are involved in stress responses (Fig. 7B, red box). Intriguingly, we found that 43 genes were senescence-specific genes, including *SAG12, SAG21*, and *SAG113* (Fig. 7B, green box), implying that those genes are common SAGs regardless of tissue-specific expression. Among the ceDEGs, we found 73 transcription factors (TF). Those genes might form a core transcriptional regulatory circuitry to coordinate the on-and-off states of gene expression during senescence. The transcription factors can be grouped into 18 families (Fig. 7C) defined from the Arabidopsis Transcription Factor Database, AtTFDB (https://agris-knowledgebase.org/AtTFDB/). The C2H2, bHLH, and AP2-EREBP families were the most abundant TF families in the ceDEGs. Many of those are involved in organ development, including *ZFP6, ZFP11*, and *JKD* (REF). Additionally, the Homeobox and MYB families were also abundant in ceDEGs. Those TFs are able to serve as “regulators” used to control their targets.

**Figure 7.**
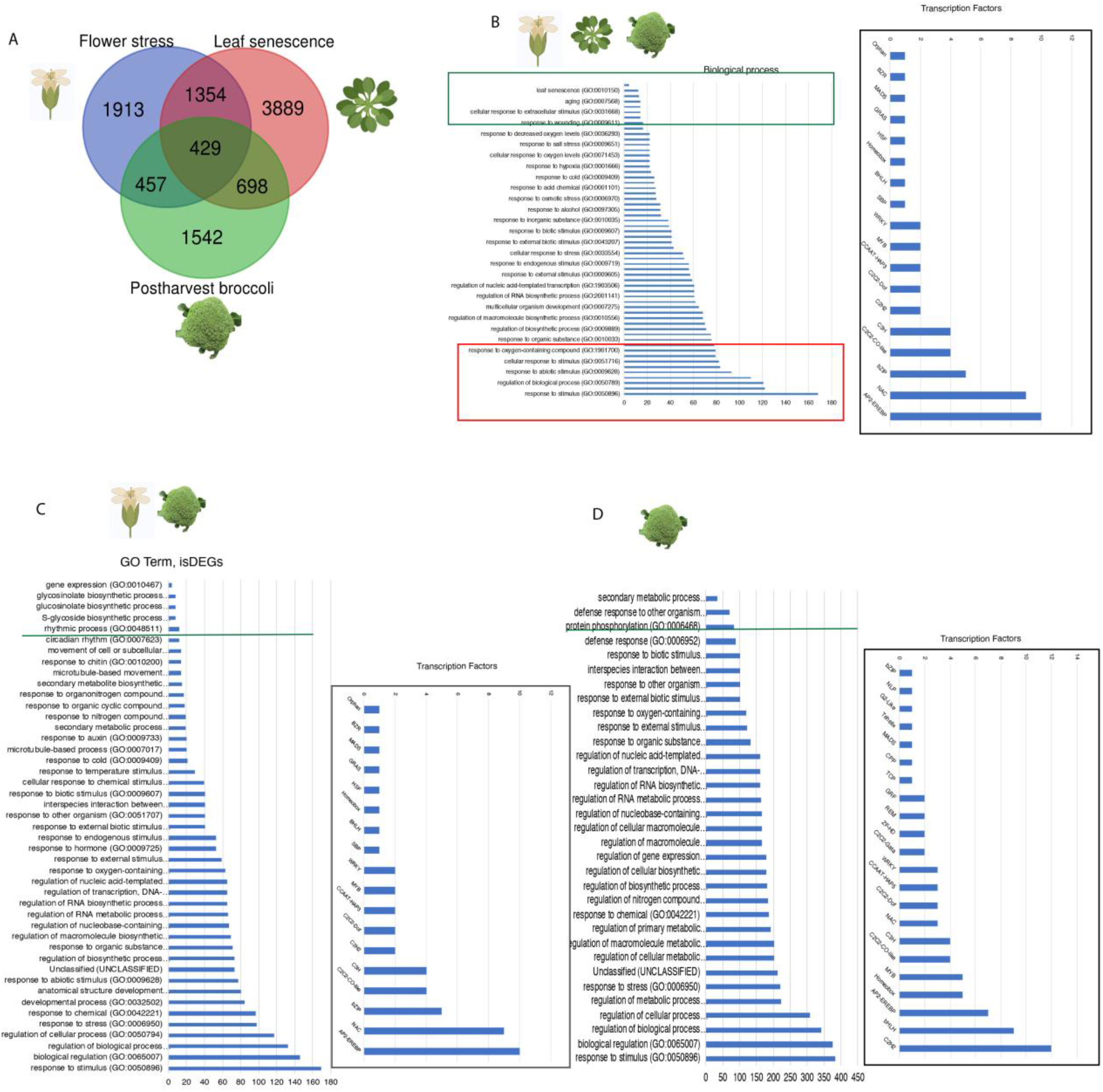
A comparison analysis among stress induced genes versus senescence-associated genes. A. Venn diagram illustrating overlapping genes among postharvest broccoli, leaf senescence (Emily, 2006) and inflorescence stress responses (Su et al. 2009). B. A list of GO terms enriched in all three datasets is shown. C. Differentially expressed transcription factors (TFs) in all three datasets, and D. TFs that were unique in broccoli.

We next leveraged the overlapping genes between postharvest broccoli and *Arabidopsis* flower senescence to identify candidate genes that are involved in tissue-specific senescence. We focused on the genes shared by those two datasets, but exclusive of the set of genes overlapped by all three datasets. Using this criteria, we defined 457 genes as inflorescence-specific DEGs (isDEGs) (Fig. 7A). Interestingly, GO analysis detected that 12 circadian clock genes were enriched in this set of genes including *CCA1, COSTANCE-Like 1* (*COL1)*, and *COL9*, which are listed in Supplementary Table 2. This is consistent with the previous observation that several SAGs such as *SAG12* and *ORE* are regulated by circadian clock-associated genes (Song et al., 2018). Additionally, five annotated senescence genes (*At1g53230, TCP3*; *At2g39730, RCA*; *At4g23810, WRKY53; At2g43570, CHI;* and *At5g5230, LTI65*) were identified in this set of genes, suggesting that those genes are flower-specific SAGs (Supplementary Table 2). Further, we identified 51 TF in this set of genes that are developmental regulators involved in inflorescence development such as *Reproductive Meristem, REM* (*REM20*), *Squamosa-Promoter binding protein-Like 3* (*SPL3*), and *Tapetal Development and Function1* (*TDF1*) (Fig. 7D, Supplementary Table 2). This suggests that those genes play general and fine-tuning roles in tissue-specific regulation of growth and stress responses.

We next examined the broccoli-specific genes using Venn diagram analysis. When we investigated the genes distinct to broccoli, we found 1542 DEGs that were not expressed in the other two datasets (Fig. 7A, green circle). Strikingly, there were 84 genes annotated in protein phosphorylation through GO terms enrichment analysis including *MAPKKK15* (*Mitogen-Activated Protein Kinase*), *CIPK22* (*CBL-Interacting Protein Kinase 22*), and *PINOID* (*PID*) (Fig. 6E, Supplementary Table 2). Another interesting group of genes within this group are the protein kinases involved in brassinosteroids (BR) perception. BRs are steroid hormones essential for plant development and stress responses. Receptor-like kinases such as *BIK1* (*Brassinosteroid-Interacting Kinase1*), *BRL1* (*BRI1-like 1*), *BIR1* (*BAK1-Inteacting Receptor-Like Kinase 1*), and *BIN2* (*Brassinosteroid-Insensitive 2*) were detected in GO term analysis (Fig. 7, Supplementary Table 2). This suggests that BRs are crucial for maintaining senescence and stress responses in postharvest broccoli. Further, we identified the TFs that are abundant in the broccoli-specific gene group (Fig. 7). We found a total of 137 TFs representing a broad range of TF families. GO enrichment analysis suggests that the TFs are classified into pathways like ‘hormone signaling’, ‘meristem development’, ‘floral whorl development’, and ‘light signaling’ (Supplementary Table 2). Taken together, these results show that our RNA-sequencing analysis enabled us to identify novel pathway regulators during postharvest senescence.

## Discussion

Fresh fruits and vegetables are invaluable for human health, but their quality deteriorates before reaching consumers during storage and distribution due to ongoing biochemical processes and compositional changes. The current lack of any objective indices for defining ‘*freshness*’ of fruits or vegetables limits our capacity to control product quality and leads to food loss and waste. Quantitative genomics analysis, such as RNA-sequencing, is a rapidly developing field that has shown the potential to expand our understanding of postharvest senescence in vegetables and fruits. Previous studies with broccoli have focused on the pigmentation changes and tissue yellowing (Luo et al. 2019). However, the mechanisms of the crosstalk between stress-responsive pathways and developmental regulation of cellular senescence that is involved in postharvest broccoli is less known. Here, we reported global transcriptional changes associated with early senescence in postharvest broccoli. We used the RNA sequencing approach to identify new genes and pathways that rapidly respond to postharvest stresses as well as to explore new players in postharvest senescence. The results increase our knowledge of early senescence and postharvest stress-response signal transduction pathways. The genomic analysis of senescence in broccoli presented here suggests that there is spatial distribution of gene senescence and that the dynamics of regulation of genes are mediated by a combination of developmental cues and environmental signals.

### Stress-responsive pathways involved in postharvest broccoli

By generating a set of transcriptomics data, we were able to quantify how transcript levels change in postharvest broccoli at room temperature and cold storage temperature. We used these two conditions because it is well-known that refrigeration slow the rate of quality deterioration (and presumably senescence processes) versus ambient conditions during postharvest handling and storage of fresh produce. We found 4279 DEGs occur in the early timepoints at room temperature, which is an estimated 9.4% of the total number of 45,758 genes in *Brassica oleracea* (Liu, et al. 2014). We found that 4143 transcripts on the fifth day of storage were DEGs, whereas there were 1408 DEGs expressed in both early and late responsive conditions. Besides, the majority of stress responsive genes, cell cycle genes, ethylene-related genes, and SAGs displaying similar expression patterns are consistently increased or decreased in both early and late postharvest response. Cellular senescence is a cell cycle arrest or cell fate transition accompanied by gene regulation of cyclin dependent kinase (Grafi, et al 2011). Interestingly, cell cycle genes include cyclins. It is known that dark-induced leaf senescence involves development of features of de-differentiating cells, which are involved in nucleolus disruption, chromatin condensation, and stem cell-like rejuvenation (Grafi et al. 2011). Considering that harvest causes premature tissue senescence, it is consistent that cell cycle genes such as cyclin genes are involved in postharvest senescence in broccoli. Additionally, we found that a range of abiotic and biotic stress-responsive genes were enriched in all timepoints in either room or low temperature conditions, including wounding, heat, oxygen, and water stresses (Fig. 2A-B).

### SAGs and developmental regulation involved in postharvest senescence

In our study, we found 43 genes that were senescence-specific genes, including several well-known SAGs such as *SAG12, SAG21*, and NAC SAGs (*NAC46, NAC87*, etc.). The NAC TFs are the positive regulators in senescence that trigger pigmentation accumulation and tissue browning (Luoni et al. 2019).

We further defined 457 genes as isDEGs. Those genes are very important for petal senescence in ornamental crops. Interestingly, there are 12 circadian clock genes among the isDEGs, including *CCA1, COSTANCE-Like 1* (*COL1*), and *COL9*. It has been shown that long-term circadian misalignment accelerated cellular senescence and led to abbreviated life in mice (Inokawa et al. 2020). In Arabidopsis, leaf senescence SAGs, including *ORE1*, have been reported to be regulated by the circadian clock regulator *CCA1* (Song et al. 2018). Another circadian clock gene, *PSEUDO-RESPONSE REGULATOR 9* (*PRR9*), positively regulated *ORESARA1* (*ORE1*) through a feed-forward regulatory loop in controlling the progress of senescence (Kim et al. 2017). Given that circadian clock genes are highly expressed in postharvest broccoli, it would be interesting to investigate their roles in postharvest responses in the future.

We also examined the tissue-specific expression of SAGs. Five annotated senescence-related genes, *At1g53230, TCP3*; At2g39730, *RCA*; *At4g23810, WRKY53*; *At2g43570*, Chitinase putative gene, *CHI;* and *At5g5230* and *LTI65* were identified in the set of isSAGs. Previous studies showed that the activities of *TCP3* were negatively regulated by miR319 in order to fine tune the onset of leaf and floral senescence (Koyama et al. 2017). *RCA* was shown to play an important role in jasmonic acid-induced leaf senescence (Shan et al. 2011). *WRKY 53* is a plant-specific transcription factor that could bind stress-related genes and SAGs and act in transcriptional feedback loops in regulating senescence signaling pathways (Miao et al. 2004). *CHI* was found to play a role in petal senescence in wallflowers (*Erysimum linifolium*; Price et al. 2008). *LTI65* is a low-temperature-induced gene that is involved in ABA-induced senescence (Shimada et al, 2020). It is important to understand how those genes are involved in postharvest senescence in broccoli; this is an area with potential for further analysis.

We also identified developmental TFs that are involved in postharvest inflorescence development, such as *Reproductive Meristem, REM* (*REM20*), *Squamosa-Promoter binding protein-Like 3* (*SPL3*), and *Tapetal Development and Function1* (*TDF1*). Those developmental factors might have a fine-tuning role in regulating the balance between growth and stress responses.

Additionally, we observed that ethylene-related genes and SAGs were continuously expressed at a high level during the tested timepoints, indicating that postharvest senescence is possibly caused by a combination of ethylene-induced and SAGs-regulated pathways.

Our genomics analysis of postharvest broccoli allows us to predict new functions of genes in senescence. One example of a senescence response involves the induction of *ZPR3*. It is well-known that *ZPR3* is a competitive inhibitor of the *HD-ZIPIII* TFs in meristem and leaf polarity development in *Arabidopsis* (Wenkel et al. 2006). Our genomics analysis and overexpression of *ZPR3* in tobacco indicated that *ZPR3* is involved in postharvest senescence in broccoli. These results are supported by studies that have elucidated the roles of *ZPR3* and *REV* (*HDZIPIII)* in controlling premature senescence by directly regulating downstream TF *WRKY53* (Xie, et al. 2014). Interestingly, we found that *SPCHLESS* (*SPCH*) (McAlister et al. 2006), a master regulator in stomatal patterning, is involved in postharvest broccoli. It would be interesting to explore how *SPCH* is involved in this network.

### Cross-regulation of hormones and stress response pathways in postharvest senescence

Crosstalk between BR and stresses such as abiotic stresses response is well known and involves key enzymes and protein kinases (Gruszka, 2018). Strikingly, there were 84 genes annotated in protein phosphorylation through GO terms enrichment analysis including *MAPKKK15* (*Mitogen-Activated Protein Kinase*), *CIPK22* (*CBL-Interacting Protein Kinase 22*), and *PINOID* (*PID*). Another interesting set of genes within this group are the protein kinases involved in brassinosteroids (BR) perception. We found receptor-like kinases such as *BIK1* (*Brassinosteroid-Interacting Kinase1*), *BRL1* (*BRI1-like 1*), *BIR1* (*BAK1-Inteacting Receptor-Like Kinase 1*), and *BIN2* (*Brassinosteroid-Insensitive 2*) are expressed during broccoli senescence (Fig. 7D and Supplementary Table 2). We found a total of 137 TFs representing a broad range of TF families that are expressed during broccoli senescence (Fig 7. Supplementary Table 2). GO enrichment analysis suggests that those TFs can be classified into pathways like hormone signaling, meristem development, floral whorl development, and light signaling, etc.

### New molecular players involved in postharvest senescence

In future work, we anticipate identifying ‘freshness indicators’ that represent distinct stages of pre-visible senescence in broccoli and other cruciferous crops and validating their precision and robustness under different conditions that occur during their postharvest storage and distribution. We can suggest that our results with broccoli can likely be extended to other related fresh cruciferous crops (i.e., cabbage, kale, arugula, cauliflower, radish, Brussels sprouts), which are mostly varieties of the same specie – *B. oleracea* – or otherwise closely related, by identifying the freshness-indicator transcripts that are most highly conserved evolutionarily among Brassicaceae species. Still broader conservation of such indicators will allow applications beyond the Brassicaceae. Tests can also be honed to include responses to diverse technologies and stresses that influence the progression of postharvest plant senescence.

The edible portion of cabbage and kale is the leaf blade, while for broccoli it is the immature inflorescence. The metabolic rate of these tissues is reflected in the respiration rate (Toivonen and Forney, 2016). After harvest, deterioration is rapid unless the product is cooled to near 0 °C; deterioration is expressed as wilting, toughening, yellowing, and flower bud opening. Visible yellowing signals the <u>final</u> stage of senescence, and is preceded by reductions in ascorbic acid, chlorophyll, soluble sugar, sucrose, starch, organic acids, and proteins (King and Morris, 1994a; Paradis et al., 1995; Pliakoni et al., 2015; Pogson and Morris, 1997; Ren et al., 2006; Starzynska et al., 2003) and increases in phenolics (Leja et al., 2001; Pliakoni et al., 2015; Starzynska et al., 2003). Broccoli inflorescences exhibit climacteric physiology during yellowing and flower opening, but ethylene production is low while broccoli remains green. However, both leaf and inflorescence tissues are extremely sensitive to ethylene as evidenced by rapid yellowing (King and Morris, 1994b; Tian et al., 1994). The role of ethylene in other (pre-visible) aspects of senescence remains unclear.

## Materials and Methods

### Collection of plant materials

Freshly grown broccoli (*Brassica oleracea*) heads from local farms in Hastings, Florida were manually harvested to avoid mechanical damage at normal commercial maturity, being fully formed, dark green, and compact. Broccoli heads of the same shape and size harvested at the same physiological age were used for the study of transcript profiling. The postharvest broccoli heads were kept under two storage conditions: cold (4 °C) and room temperature (RT, 25 °C). The heads were stored in a cold room (4–5 °C, darkness) for the cold treatment, or in a plant growth chamber at 25°C (RT) under the same conditions. Three biological replicates of broccoli florets (100 mg) removed from the stored heads were used for RNA sequencing and proteomics. Floret samples were collected from the broccoli bunches on days 1, 3, and 5 and wrapped in aluminum foil. The samples were immediately frozen in liquid nitrogen and then stored at -80 °C for later use.

### Chlorophyll assay

Broccoli heads that were stored at RT (25 °C) or 4 °C were measured for the total chlorophyll content on days 1, 3, 5, 7, and 9. A total of 400 mg of floret tissues were ground under liquid nitrogen using a mortar and pestle and the resulting powder was dissolved in 5 ml of 80% acetone (v/v). The dissolved mixture was then centrifuged at 8000×g_n_ for 15 minutes at 4 °C. The supernatant was used for measuring the absorbance at 645 nm and 663 nm and the total chlorophyl content was calculated using the formula, Chl a (mg g-1) = [(12.7 × A663) - (2.6 × A645)] × ml acetone / mg leaf tissue Chl b (mg g-1) = [(22.9 × A645) - (4.68 × A663)] × ml acetone / mg leaf tissue Total Chl = Chl a + Chl b. Four biological replicates were measured on each day of sampling. Total chlorophyll content was expressed as milligrams of total chlorophyll per gram of fresh tissue taken for sampling.

### Water loss measurement

Water loss was monitored by measuring the fresh weight of the broccoli heads using four biological replicates from each time point. Five time points were chosen respectively on day 1, day 3, day 5, day 7, and day 9 when broccoli heads were stored at RT or at 4 °C. Water loss was calculated by averaging the biological replicates from each time point.

### RNA Extraction

Broccoli floret tissue samples from days 1, 3, and 5 were chosen for the total transcriptome analysis. There were 18 samples in total, 3 biological? replicates each at days 1, 3, and 5 from two storage conditions-25 °C and 4 °C. Total RNA was isolated from broccoli floret tissue using Trizol (Ambion, Life Technologies) following the manufacturer’s protocol. Extracted RNA was subjected to DNase treatment (Turbo DNA free, Thermo-Fisher). The 260/280 nm and 260/230 nm absorption ratios were further tested on Nanodrop (Applied Biosystems) to investigate for their concentration and purity. Quality of total RNA for each sample was determined using an Agilent 2100 Bioanalyzer (Agilent Technologies, Santa Clara, CA, USA) at the University of Florida Interdisciplinary Center for Biotechnology Research (ICBR), Gene Expression and Genotyping Core Lab.

### RNA library preparation and RNA-sequencing

RNA-seq libraries were constructed at the ICBR Gene Expression & Genotyping Core Lab using NEBNext® Ultra™ Directional RNA Library Prep Kit for Illumina (NEB, USA) following the manufacturer’s recommendations. Basically, 1000 ng samples of high quality total RNA (RIN≥ 7) was used for mRNA isolation using NEBNext Ploy(A) mRNA Magnetic Isolation module (New England Biolabs, catalog # E7490), followed by RNA library construction with NEBNext Ultra II Directional Lib Prep (New England Biolabs, catalog #E7760) according to the manufacturer’s user guide. Briefly, RNA was fragmented in NEBNext First Strand Synthesis Buffer by heating at 94 °C for desired time. This step was followed by first strand cDNA synthesis using reverse transcriptase and oligo dT primers. Synthesis of ds cDNA was performed using the 2nd strand master mix provided in the kit, followed by end-repair and adaptor ligation. Finally, the library was enriched (each library has a unique barcode) by 8 cycles of amplification, and purified by Agencourt AMPure beads (Beckman Coulter, catalog # A63881). Finally, the 18 individual libraries were pooled using equimolar amounts and sequenced by Illumina NovaSeq 2×150 cycles run for a total of 0.5 lane (Illumina Inc., CA, U.S.)

### Data analysis

FastQC was used to evaluate the RNA sequencing quality and then Trimmomatic was used to trim low quality reads with phred score < 20 within windows of 4 base pairs (Folger et al., 2014). The reads after trimming with length < 80 were removed from downstream analysis. The remaining high quality reads were aligned to a *Brassica oleracea* reference genome (Kim et al., 2014) (https://www.ncbi.nlm.nih.gov/genome?LinkName=nuccore_genome&from_uid=601087403) using Bowtie with default parameters. Gene expressions at the gene level and isoform level were obtained using RSEM (Li and Dewey, 2011). EdgeR was used to perform differential analysis to calculate log2 fold change between each two comparisons (Chen et al., 2014).). PCA was performed to identify outlier samples. No obvious outlier samples were found.

### Quantitative PCR (QRT-PCR) analysis

Transcript level studies were performed to validate the differentially expressed genes observed during the RNA-sequencing analysis from the postharvest broccoli stored at day 1, 3, and 5. Relative transcript expression was investigated by using three technical replicates from each of the biological replicates from these three time points. Actin 2 gene was used as an internal reference control and the DEGs primers are included in Supplementary Table 3. Each reaction was 12 µl using SYBR green reagents from a Power up SYBR green kit (Thermo-Fisher, USA). Each gene was amplified in three separate reactions using a Thermofisher Applied biosystems thermocycler with the following conditions: 95°C for 10 min, 45 cycles for 95°C for 30 sec, 60°C for 30 sec. Relative gene expression was calculated by the ∆∆CT method. Primer sequences were listed in Supplementary Table 3.

### Senescence assay in tobacco

Transient gene expression in tobacco plants (*Nicotiana benthamiana*) was investigated to determine the subcellular location for the selected DEGs observed in our sequencing data. The selected candidate genes were fused to pMDC43 vector (https://www.arabidopsis.org/servlets/TairObject?type=vector&id=501100109) which contained a 35S promoter and GFP tag. GV3101 *Agrobacterium* strain was used to infiltrate the tumor-inducing plasmid into the 28-day-old tobacco epidermal cells. We inoculated one single colony of 8 agrobacterium transformants (along with one empty vector as control) in 5 ml YEB with appropriate antibiotics rifampicin (50 μg/ml) and kanamycin (50μg/ml). 500 μl of the primary overnight culture was used for subculturing during secondary inoculation using the same selection antibiotic. During the transient transformation in tobacco, 100 ml of re-suspension media MES (10 mM) and magnesium chloride (10 mM) with 100 μM acetosyringone was used to dissolve the pellet from the secondary culture (bacteria from the secondary culture were precipitated at 5,000 x g for 15 min). The OD was adjusted to 0.5 at A600 in the secondary overnight culture and the infiltration media. The tobacco leaves were infiltrated using a 5-ml syringe on the ventral surface of the leaf using counter pressure by a finger on the dorsal side. Successful infiltration could be easily observed through the wet surface of the leaves on the day of injection. The transient tobacco assays have been repeated for three times with four biological replicates for each agrobacteria culture containing the pMDC43 construct. Images from the tobacco leaves were taken immediately and at the interval of every 3 days. RNA was extracted from the control and those inoculated leaves followed by cDNA synthesis and qPCR experiment as described above.

## Acknowledgements

We thank extension specialist Prissy Fletcher provided broccoli samples from local farm in Florida. We also thank UF ICBR Gene Expression & Genotyping and Bioinformatic facilities conducted RNA-sequencing and raw data analysis.

## Figure legends

**Supplementary Figure 1. Validation of genes involved in senescence through tobacco transient assay. A**. Representative images of control tobacco leaves injected with Overexpressing *SSP (Senescence Suppressed Phosphatase* (*pMDC43::SSP*) in tobacco leaves caused accelerated tissue senescence. B. Overexpressing of *S-adenosyl-L-methionine-dependent methyltransferases (S-ALM)* (*pMDC43::S-ALM)* in tobacco leaves displayed progressive senescence after injection. C. Overexpression of NAC46 (*pMDC43::NAC46*).

**Supplementary Figure 1. Test candidate genes involved in senescence through *Arabidopsis* mutant analysis**. Selected DEGs of *Arabidopsis* homologues and their T-DNA insertion mutants. A-B. *SALK072093C*, a loss-of-function of At5g00875 showed delayed maturation and senescence phenotypes. C-D. *SALK027396C*, a loss-of-function of *s-alm*. E-F. *SALK203275C*, a loss-of-function of *zpr3*, showed delayed senescence.

**Supplementary Table1. Comparative analysis of Arabidopsis and broccoli senescence gene-regulatory network (GRN)**

**Supplementary Table2. GO and Transcription Factor enrichment**.

**Supplementary Table3. Primer sequences for qPCR analysis**

## References

Ahlawat, Y. and Liu, T., 2021. Varied Expression of Senescence-Associated and Ethylene-Related Genes during Postharvest Storage of Brassica Vegetables. International Journal of Molecular Sciences,22(2), p.839.doi: 10.3390/ijms22020839

Ashby, E. (1950). Leaf Morphology and Physiological Age. Science Progress (1933-), 38(152), 678–685. Retrieved February 3, 2021, doi: http://www.jstor.org/stable/43413975

Breeze, E. et al. (2011) ‘High-Resolution Temporal Profiling of Transcripts during Arabidopsis Leaf Senescence Reveals a Distinct Chronology of Processes and Regulation’, The Plant Cell, 23(3), pp. 873–894. doi: 10.1105/tpc.111.083345.

Bolger, A.M., Lohse, M. and Usadel, B., 2014. Trimmomatic: a flexible trimmer for Illumina sequence data. Bioinformatics, 30(15), pp.2114–2120.

Chen, Y., Lun, A.T. and Smyth, G.K., 2014. Differential expression analysis of complex RNA-seq experiments using edgeR. Statistical analysis of next generation sequencing data, pp.51–74.

Chen, Y.-T., Chen, L.-F. O. and Shaw, J.-F. (2008) ‘Senescence-associated genes in harvested broccoli florets’, Plant Science, 175(1–2), pp.137–144. doi: 10.1016/j.plantsci.2008.03.007.

Cliff, M. A. and Toivonen, P. M. A. (2017) ‘Sensory and quality characteristics of “Ambrosia” apples in relation to harvest maturity for fruit stored up to eight months’, Postharvest Biology and Technology, 132, pp.145–153. doi: 10.1016/j.postharvbio.2017.05.015.

Coupe, S. A. et al. (2003) ‘Analysis of acid invertase gene expression during the senescence of broccoli florets’, Postharvest Biology and Technology, 28(1), pp.27–37. doi: 10.1016/S0925-5214(02)00126-6.

Damari-Weissler, H. et al. (2009) ‘LeFRK2 is required for phloem and xylem differentiation and the transport of both sugar and water’, Planta, 230(4), pp.795–805. doi: 10.1007/s00425-009-0985-4.

Eason, J. R. et al. (2005) ‘Suppression of the cysteine protease, aleurain, delays floret senescence in Brassica oleracea’, Plant Molecular Biology, 57(5), pp.645–657. doi: 10.1007/s11103-005-0999-7.

Eason, J. R. et al. (2014) ‘Overexpression of the protease inhibitor BoCPI-1 in broccoli delays chlorophyll loss after harvest and causes down-regulation of cysteine protease gene expression’, Postharvest Biology and Technology, 97, pp.23–31. doi: 10.1016/j.postharvbio.2014.06.006.

Gapper, N. E. et al. (2005) ‘Regulation of Harvest-induced Senescence in Broccoli (Brassica oleracea var. italica) by Cytokinin, Ethylene, and Sucrose’, Journal of Plant Growth Regulation, 24(3), pp.153–165. doi: 10.1007/s00344-005-0028-8.

Gonzalez, N. and Botella, J. R. (2003) ‘Characterisation of three ACC synthase gene family members during post-harvest-induced senescence in broccoli (Brassica oleracea L. var. italica)’, Journal of Plant Biology, 46(4), pp.223–230. doi: 10.1007/BF03030368.

Grafi, G. et al. (2011) ‘Plant response to stress meets dedifferentiation’, Planta, 233(3), pp.433– 438. doi: 10.1007/s00425-011-1366-3.

Humbeck, K. (2014) ‘Senescence in Plants’, Journal of Plant Growth Regulation, 33(1), pp.1–3. doi: 10.1007/s00344-013-9397-6.

Inokawa, H. et al. (2020) ‘Chronic circadian misalignment accelerates immune senescence and abbreviates lifespan in mice’, Scientific Reports, 10(1), p.2569. doi: 10.1038/s41598-020-59541-y.

Kamachi, K. et al. (1991) ‘A Role for Glutamine Synthetase in the Remobilization of Leaf Nitrogen during Natural Senescence in Rice Leaves’, Plant Physiology, 96(2), pp.411–417. doi: 10.1104/pp.96.2.411.

Kim, H. et al. (2018) ‘Circadian control of ORE1 by PRR9 positively regulates leaf senescence in Arabidopsis’, Proceedings of the National Academy of Sciences, 115(33), pp.8448–8453. doi: 10.1073/pnas.1722407115.

Kim, H.A., Lim, C.J., Kim, S., Choe, J.K., Jo, S.H., Baek, N. and Kwon, S.Y., 2014. High-throughput sequencing and de novo assembly of Brassica oleracea var. Capitata L. for transcriptome analysis. PLoS One, 9(3), p.e92087.

King, G. A. and Morris, S. C. (1994) ‘Physiological Changes of Broccoli during Early Postharvest Senescence and through the Preharvest-Postharvest Continuum’, Journal of the American Society for Horticultural Science, 119(2), pp.270–275. doi: 10.21273/JASHS.119.2.270.

Koyama, T., Sato, F. and Ohme-Takagi, M. (2017) ‘Roles of miR319 and TCP Transcription Factors in Leaf Development’, Plant Physiology, 175(2), pp.874–885. doi: 10.1104/pp.17.00732.

Leja, M. et al. (2001) ‘Antioxidant ability of broccoli flower buds during short-term storage’, Food Chemistry, 72(2), pp.219–222. doi: 10.1016/S0308-8146(00)00224-7.

Langmead, B. and Salzberg, S.L., 2012. Fast gapped-read alignment with Bowtie 2. Nature methods, 9(4), p.357

Li, B. and Dewey, C.N., 2011. RSEM: accurate transcript quantification from RNA-Seq data with or without a reference genome. BMC bioinformatics, 12(1), pp.1–16.

Li, Z. et al. (2012) ‘Gene Network Analysis and Functional Studies of Senescence-associated Genes Reveal Novel Regulators of Arabidopsis Leaf SenescenceF’, Journal of Integrative Plant Biology, 54(8), pp.526–539. doi: 10.1111/j.1744-7909.2012.01136.x.

Luo, F. et al. (2019) ‘Chlorophyll degradation and carotenoid biosynthetic pathways: Gene expression and pigment content in broccoli during yellowing’, Food Chemistry, 297, p.124964. doi: 10.1016/j.foodchem.2019.124964.

Nooden, L. D., Guiamet, J. J. and John, I. (1997) ‘Senescence mechanisms’, Physiologia Plantarum, 101(4), pp.746–753. doi: 10.1111/j.1399-3054.1997.tb01059.x.

Page, T., Griffiths, G. and Buchanan-Wollaston, V. (2001) ‘Molecular and Biochemical Characterization of Postharvest Senescence in Broccoli’, Plant Physiology, 125(2), pp.718–727. doi: 10.1104/pp.125.2.718.

Péneau, S. et al. (2006a) ‘Importance and consumer perception of freshness of apples’, Food Quality and Preference, 17(1–2), pp.9–19. doi: 10.1016/j.foodqual.2005.05.002.

Péneau, S. et al. (2006b) ‘Importance and consumer perception of freshness of apples’, Food Quality and Preference, 17(1–2), pp.9–19. doi: 10.1016/j.foodqual.2005.05.002.

Pogson, B. J. and Morris, S. C. (1997) ‘Consequences of Cool Storage of Broccoli on Physiological and Biochemical Changes and Subsequent Senescence at 20 °C’, Journal of the American Society for Horticultural Science, 122(4), pp.553–558. doi: 10.21273/JASHS.122.4.553.

Price, A. M. et al. (2008) ‘A Comparison of Leaf and Petal Senescence in Wallflower Reveals Common and Distinct Patterns of Gene Expression and Physiology’, Plant Physiology, 147(4), pp.1898–1912. doi: 10.1104/pp.108.120402.

Rachappanavar, V. et al. (2020) PLANT HORMONE-MEDIATED REGULATION OF STRESS RESPONSES IN FRUIT CROPS-A REVIEW. preprint. Preprints. doi: 10.22541/au.160466945.56576285/v1.

Ragaert, P. et al. (2004) ‘Consumer perception and choice of minimally processed vegetables and packaged fruits’, Food Quality and Preference, 15(3), pp.259–270. doi: 10.1016/S0950-3293(03)00066-1.

Ren, K. et al. (2006) ‘KINETIC MODELINGS OF BROCCOLI COLOR CHANGES DURING CHILLED STORAGE: KINETIC MODELS OF COLOR CHANGES OF STORED BROCCOLI’, Journal of Food Processing and Preservation, 30(2), pp.180–193. doi: 10.1111/j.1745-4549.2006.00058.x.

Reyes Jara, A. M. et al. (2019) ‘Effects of hormonal and physical treatments on the expression of a putative chlorophyll b reductase gene (BoNYC1) during postharvest senescence of broccoli’, Postharvest Biology and Technology, 147, pp.107–112. doi: 10.1016/j.postharvbio.2018.09.010.

Shan, X. et al. (2011) ‘The Role of Arabidopsis Rubisco Activase in Jasmonate-Induced Leaf Senescence’, Plant Physiology, 155(2), pp.751–764. doi: 10.1104/pp.110.166595.

Shimada, T. L. et al. (2020) ‘Excess sterols disrupt plant cellular activity by inducing stress-responsive gene expression’, Journal of Plant Research, 133(3), pp.383–392. doi: 10.1007/s10265-020-01181-4.

Song, Y. et al. (2018) ‘CIRCADIAN CLOCK-ASSOCIATED 1 Inhibits Leaf Senescence in Arabidopsis’, Frontiers in Plant Science, 9, p.280. doi: 10.3389/fpls.2018.00280.

Starzynska, A., Leja, M. and Mareczek, A. (2003) ‘Physiological changes in the antioxidant system of broccoli flower buds senescing during short-term storage, related to temperature and packaging’, Plant Science, 165(6), pp.1387–1395. doi: 10.1016/j.plantsci.2003.07.004.

Wenkel, S. et al. (2006) ‘CONSTANS and the CCAAT Box Binding Complex Share a Functionally Important Domain and Interact to Regulate Flowering of Arabidopsis’, The Plant Cell, 18(11), pp.2971–2984. doi: 10.1105/tpc.106.043299.

Xie, Y. et al. (2014) ‘REVOLUTA and WRKY53 connect early and late leaf development in Arabidopsis’, Development, 141(24), pp.4772–4783. doi: 10.1242/dev.117689.

